# EcoEnamel: Development of a Gelatin-Pectin Film for *S. mutans* Inhibition and Enamel Preservation in an In Vitro Model

**DOI:** 10.64898/2026.08.18.745620

**Authors:** Gabrielle Javelona, Jacob Merle

**Affiliations:** Science Department, Vanguard High School, Ocala, Florida; Science Department, The Cornerstone School, Ocala, Florida

**Keywords:** Biofilm, Hydroxyapatite, Caries, Antimicrobial, Remineralization

## Abstract

Rinsing-dependent dental hygiene presents a significant public health challenge in water-scarce environments. This study investigated combinations of xylitol (Xyl), chitosan (Chi), glycyrrhizin (Gly), epigallocatechin gallate (EGCG), dicalcium phosphate (DCP), and nano-hydroxyapatite (nHA) on the primary bacteria behind dental caries, *S. mutans*. These combinations were assessed for markers of dental caries by biofilm reduction, bacterial killing, and acid buffering against *S. mutans* when applied to an in vitro simulated enamel model using glass bead surfaces for biofilm formation, and gene expression was subsequently examined via RT-qPCR. Separately, mineral retention was also quantified. The EGCG-DCP-Xyl film demonstrated the highest overall efficacy, achieving a significant reduction in biofilm concentration compared to the untreated control and performing similarly in magnitude to the positive toothpaste control. Dead fluorescence staining confirmed that the EGCG-DCP-Xyl film induced the highest rate of non-viable cells, followed by the Chi-Gly film and the Gly-Xyl film. During 10-day pH cycling, the EGCG-DCP-Xyl and DCP-Xyl formulations buffered pH the most, consistently maintaining mean pH levels safely above the demineralization threshold of pH 5.5. The EGCG-DCP-Xyl also optimized mineral stability with the highest retained calcium concentration, significantly outperforming the Chi-Xyl film. At the transcript level, the EGCG-DCP-Xyl film induced substantial downregulation of key virulence genes, yielding decreases in expression for glucosyltransferase B (gtfB), associated with biofilm synthesis, collagen-binding protein (cnm), associated with tissue invasion, and lactate dehydrogenase (ldh), associated with lactic acid production, compared to the untreated control, with effects comparable in magnitude to the positive toothpaste control. This research suggests that targeting bacterial pathways and mineral loss through a portable film may have potential for preventing dental caries, especially in environments where water is limited. However, additional studies are necessary to evaluate real-world effectiveness.

## INTRODUCTION

Dental caries is the most common chronic disease globally, affecting roughly 2.5 billion people (1). The primary bacterium behind this decay is the Gram-positive Streptococcus mutans (2). If left untreated, dental caries can progress to more severe oral infections or broader systemic health risks. (3).

Currently, dentifrices (toothpastes) are the primary defense against dental caries, but these require rinsing with clean water. In water-scarce environments, the absence of rinsing infrastructure introduces a significant public health challenge: individuals must either forgo oral hygiene entirely or swallow concentrated dentifrice residue (1). For children younger than eight years of age, permanent teeth undergo amelogenesis, and repeatedly ingesting this residue during enamel development can alter the minerals in the enamel and increase the risk of dental fluorosis, especially since swallowing behavior is not adequately controlled at these ages (2). This is why pediatric dental guidelines emphasize supervision and limited toothpaste quantities for young children, however this can be difficult to enforce in rinsing-inadequate environments.

Independent of fluorosis concerns, conventional toothpaste use generally relies on access to clean water for rinsing after application. For populations living in water-scarce environments, maintaining rinsing-based oral hygiene practices may be more difficult due to limitations in water availability and sanitation infrastructure. Approximately 2.2 billion people worldwide lack safely managed drinking water services, highlighting the need for alternative oral health interventions that require little or no water during use (1).

To address this limitation, a water-free, dissolvable, and ingestible film was developed to deliver therapeutic agents directly to the tooth surface. Alternative, non-toxic, edible combinations of ingredients categorized as Generally Recognized as Safe (GRAS) by the FDA were selected based on existing literature (4). Ingredients were selected based on their established use in food, pharmaceutical, and oral-care products, as well as their favorable safety profiles at the concentrations employed in this study. However, “safer” does not mean biologically inert or universally non-toxic. Each compound may produce adverse effects at sufficiently high doses or with chronic exposure. Therefore, these ingredients are better described as candidate alternatives with comparatively favorable expected safety profiles at low, localized oral doses, rather than as categorically non-toxic. (see Table 1 for LD50 levels)

**Table 1:**
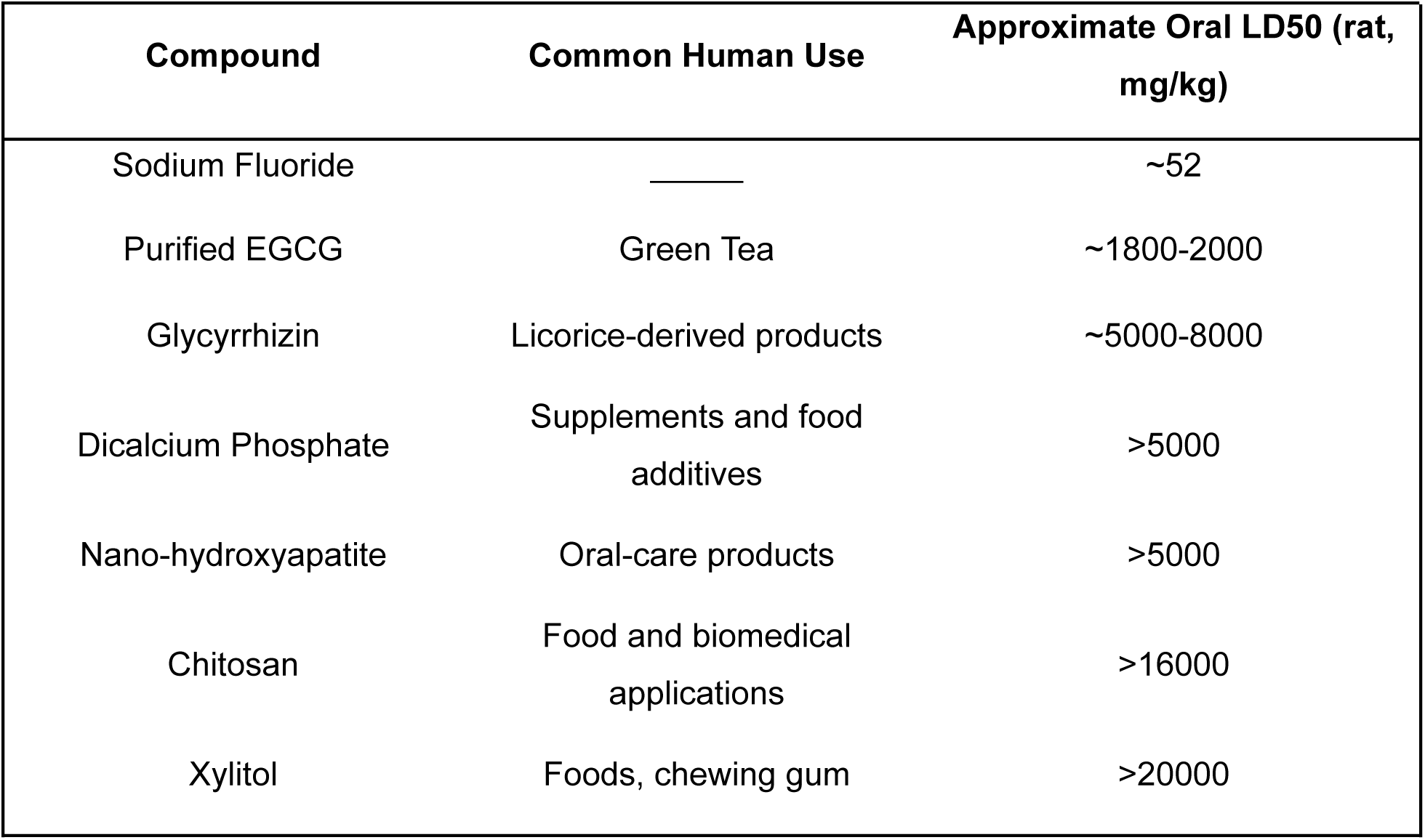
Representative oral acute toxicity values and safety considerations for fluoride and the various formula ingredients. (9, 10, 11, 14, 16, 17) LD_50_ values are reported from published animal studies and are provided to illustrate relative acute toxicity. These values should not be interpreted as indicators of chronic safety, acceptable daily intake, or clinical risk.

| Compound | Common Human Use | Approximate Oral LD50 (rat, mg/kg) |
| --- | --- | --- |
| Sodium Fluoride | _____ | ~52 |
| Purified EGCG | Green Tea | ~1800-2000 |
| Glycyrrhizin | Licorice-derived products | ~5000-8000 |
| Dicalcium Phosphate | Supplements and food additives | >5000 |
| Nano-hydroxyapatite | Oral-care products | >5000 |
| Chitosan | Food and biomedical applications | >16000 |
| Xylitol | Foods, chewing gum | >20000 |

The ingredients include xylitol for acid reduction (9), licorice and green tea extract (epigallocatechin gallate, EGCG) as allosteric inhibitors to *S. mutans* (18), chitosan to disrupt bacterial cell walls via positive charge interactions (11), and dicalcium phosphate (DCP) and nano-hydroxyapatite (nHA) to promote mineral retention (15).

*S. mutans* is a widely used model used to study the progression of dental caries due to its ability to synthesize extracellular polysaccharides and to generate an acidic microenvironment (2). When these bacteria are disturbed by the active compounds in the film delivery system, their metabolic pathways are significantly altered. For example, chitosan has a high polycationic charge, which allows it to positively bind to the negatively charged *S. mutans* cell membrane. This causes intracellular component leakage, and eventually cell lysis (Figure 1, Source 11). Furthermore, xylitol specifically inhibits the lactate dehydrogenase (*ldh)* pathway.

**Figure 1:**
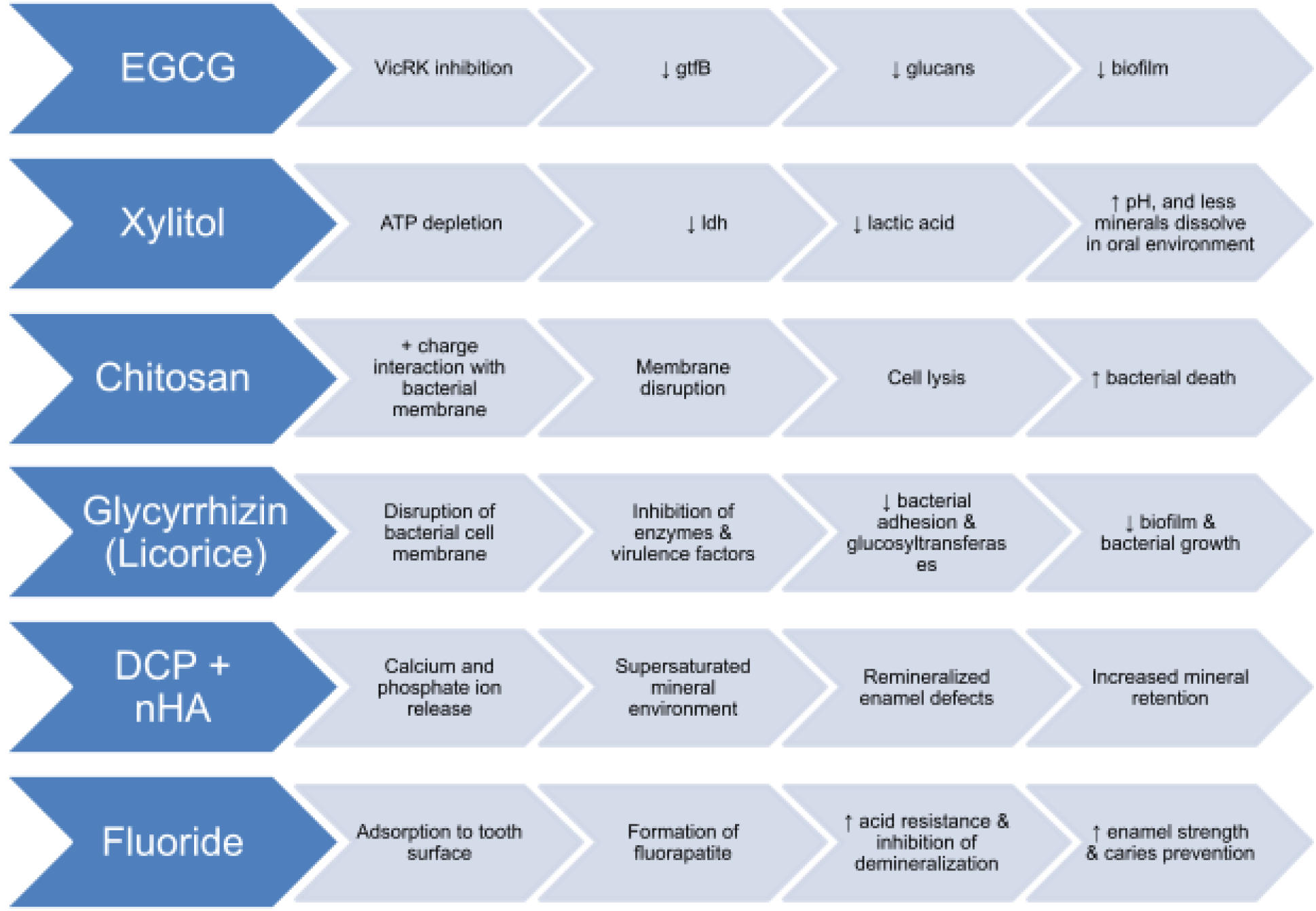
Proposed mechanisms by which the various active ingredients in the film may influence dental caries-associated processes. Antimicrobial compounds (EGCG, glycyrrhizin, chitosan, and xylitol) have been reported to reduce Streptococcus mutans growth, virulence, biofilm formation, and acid production (9, 10, 11, 18, 19), while DCP and nHA may promote mineral retention and remineralization (12, 15, 16, 17). These are provided compared to fluoride (9). These mechanisms are hypothesized based on previous literature and are presented as a conceptual framework within the present study.

In *S. mutans,* the LDH pathway is the critical final step of anaerobic glycolysis. It catalyzes the reduction of pyruvate to lactic acid, regenerating the NAD⁺ needed for continuous ATP production (2). The resulting secretion of lactic acid rapidly lowers the environmental pH. This local acidification creates the acidic microenvironment responsible for tooth enamel demineralization and dental caries progression (Figure 1).

Xylitol forces *S. mutans* to expend ATP trying to expel it, which produces a negative feedback loop, leading to the downregulation of the entire *ldh* pathway (9). Licorice has the active compound glycyrrhizin, an allosteric inhibitor that interferes with protein synthesis (8). Green tea has the active compound epigallocatechin gallate (EGCG), which is a more effective allosteric inhibitor (14). EGCG targets the virulence control response regulator and histidine kinase (VicRK) signal transduction system, which regulates the *gtfB* gene.

The VicRK two-component system in *S. mutans* regulates virulence by sensing environmental stimuli to activate the response regulator VicR. Activated VicR binds directly to the gtfB promoter, upregulating transcription of glucosyltransferase B (20). This enzyme synthesizes water-insoluble glucans, which build the extracellular polysaccharide (EPS) matrix necessary for future biofilm formation and bacterial adherence (Figure 1). By inhibiting this system, EGCG prevents the bacteria from sensing and responding to their environment, shutting down the production of those adhesive glucans and therefore also shutting down overall biofilm structure (14).

Tooth enamel is made of a mineral called hydroxyapatite, which is a combination of calcium and phosphate. Dicalcium phosphate (DCP) is known to restore these minerals to the simulated tooth surface (16), and nano-hydroxyapatite incorporated in all test formulas also plugs any micropores and incipient lesions created by *S. mutans* acid challenges, restoring the natural tooth structure (17).

The ingredients in the different film test formulas have been tested through various in vivo studies, with results reflecting high therapeutic potential. One study found that nano-hydroxyapatite (nHA) particles at concentrations of 5-10% can fill in enamel lesions and may even restore surface microhardness in bovine enamel models (15). Similarly, a high-sucrose diet supplemented with green tea (EGCG) has been shown to reduce the incidence of dental caries by nearly 40% in rodent models (18).

Since these individual compounds target the three different pillars of *S. mutans* activity and also address enamel demineralization, they may be even more effective in combination. Therefore, it was hypothesized that a EGCG-DCP-Xyl film would show the strongest effects across all measured endpoints, including biofilm levels, bacterial viability, pH stability, mineral retention, and gene expression.

## MATERIALS AND METHODS

Two distinct in vitro models were utilized to independently simulate the physical and chemical properties of tooth enamel. First, a surface-based microbiology model utilizing inert glass beads was established to provide a standardized, uniform substrate for *S. mutans* biofilm attachment and viability analysis. Separately, a chemical mineralization model utilizing calcium tablets was employed to quantify enamel mineral retention and acid buffering dynamics during challenge phases. This dual-model framework ensured that surface-level bacterial colonization and deeper demineralization kinetics were evaluated independently. Three experimental phases were developed to evaluate the multi-targeted efficacy of the formulations relative to standard dentifrice controls. First, Colony Forming Units (CFU) counts were used to quantify biofilm burden and measure the capacity of each formula to disrupt bacterial accumulation; a higher concentration of culturable cells indicates greater overall biofilm development on the substrate (5). Second, dead fluorescence staining measured how well each formula would kill the bacteria living on the tooth surface (6), and third, pH cycling was used to measure how well each formula would neutralize the acids produced by *S. mutans* in the simulated oral cavity (7). The mineral retention of each formula was measured using a calcium assay. Lastly, RT-qPCR was used to analyze the gene expression in *S. mutans* after being exposed to the treatments for genetic context.

### Preparation of Test Films

The film base was prepared by heating 100 mL of distilled water to 50°C. Gelatin (2.5 g), pectin (1 g), glycerin (1 mL), sodium benzoate (0.15 g), and nano-hydroxyapatite (1.5 g) were dissolved and incorporated using a magnetic stirrer. Five distinct formulations were then synthesized by incorporating varying active ingredients (Table 2). For formulas containing chitosan, 0.5 g was pre-dissolved in 10 mL of 1% acetic acid prior to integration. Once homogenous, the mixtures were poured into 5×5 cm templates made with parchment paper and dehydrated for 13 hours. Final films were sectioned into 3×3 mm squares for experimental application.

**Table 2:** Composition of Experimental Treatment Formulations (Weight By Volume): Descriptions of the active ingredient combinations for each experimental formula (1–5) are provided here. Formulas were standardized to ensure consistent delivery of ingredients, with concentrations provided adjacent to each ingredient.

| 1 | 2 | 3 | 4 | 5 |
| --- | --- | --- | --- | --- |
| Chi-Xyl | Chi-Gly | DCP-Xyl | Gly-Xyl | EGCG-DCP-Xyl |
| nHA (1.5%) | nHA (1.5%) | nHA (1.5%) | nHA (1.5%) | nHA (1.5%) |
| Xylitol (1.5%) | Licorice (1%) | DCP (1.5%) | Licorice (1%) | Xylitol (1.5%) |
| Chitosan (0.5%) | Chitosan (0.5%) | Xylitol (1.5%) | Xylitol (1.5%) | Green Tea |
|  |  |  |  | DCP (1.5%) |

### Bacterial Culture and Inoculum Preparation

Streptococcus mutans was cultured on nutrient agar and incubated at 37°C. Liquid cultures were prepared by inoculating a single colony into TSB liquid media and grown to a standardized optical density (OD600) of 0.10, measured with a spectrophotometer. A simulated saliva solution (mucin saliva) was prepared containing mucin (2.4 g/L), urea (0.16 g/L), and essential salts to mimic the oral environment.

**Figure 2A:**
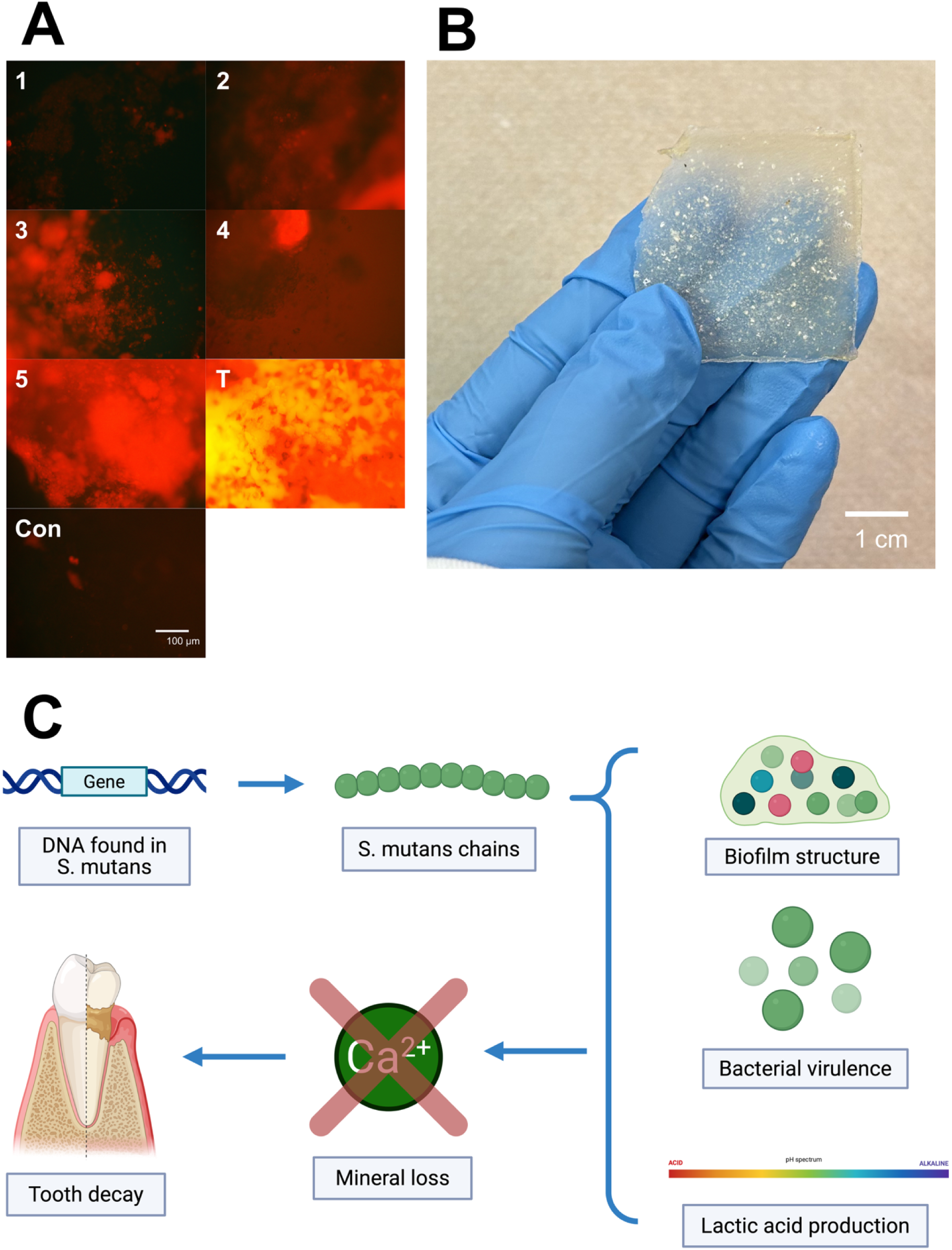
**Images showing the presence and distribution of nonviable, stained bacteria:** A fixable dye was applied to highlight unviable localized bacterial clusters within the biofilm matrix (n=5). Labels 1-5 represent the experimental groups shown in a single representative trial (Trial 1 of 5). T represents toothpaste, and Con represents the untreated control. The percentages of dead cells in these images were as follows: <10%, 37%, 67%, 42%, 55%, 75%, 79%. One-way ANOVA followed by Tukey’s HSD post-hoc tests (p<0.05). **Figure 2B: The EcoEnamel Film:** A visual of the experimental film delivery system used in the study. The material is translucent and embedded with therapeutic ingredients. **Figure 2C: *S. mutans* to Tooth Decay Overview:** A flow diagram illustrating the process being inhibited: The film treatments inhibit the gene expression of *S. mutans*, which affects their chain communication. This yields less biofilm structures, decreased virulence, and less lactic acid production, preserving Ca and reducing tooth decay.

### Biofilm Viability Analysis (CFU Counts)

To simulate dental pellicle formation, 140 glass beads were placed individually into 96-well plates and submerged in 0.1 mL of simulated mucin saliva for 2 hours. Wells were then aspirated and inoculated with 0.2 mL of standardized *S. mutans* culture. EcoEnamel films (3×3 mm) were applied to the treatment wells, while Sensodyne Pronamel served as a positive control and distilled water as a negative control. After 24 hours of incubation, biofilms were dislodged, serially diluted, and 0.1 mL plated on nutrient agar. Colony Forming Units (CFU) were quantified after a further 24 hours of incubation. These were translated into their original concentration using the following formula:

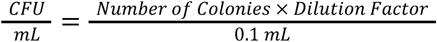

### Fluorescence Microscopy for Cell Mortality

Bacterial killing efficiency was evaluated using a fixable red dead cell stain kit (Invitrogen). This dye enters cells through compromised cell membranes and reacts with intracellular proteins, displaying red fluorescence. The samples were imaged through fluorescent microscopy and quantified using ImageJ software to determine the mean percentage of dead cells.

After treatment, wells were washed with Phosphate-Buffered Saline (PBS) and incubated with fluorescent dye for 15 minutes in the dark. Images were captured using an OMAX fluorescent microscope under a blue laser (570 nm). The ratio of non-viable (red) to total cells was quantified using ImageJ software.

### pH Cycling

A 10-day in vitro pH cycling model was established using calcium tablets to simulate enamel demineralization. Each cycle consisted of an acid challenge (Sprite) for 2 hours, followed by an alkaline challenge (skim milk) for 22 hours. pH measurements were recorded using a Beetrode micro-pH electrode at 0, 1, and 2 hours after acids, and 0. 16, and 22 hours after alkaline solutions. The electrode was calibrated daily using a 3-point curve (pH 4.0, 7.0, and 10.0) with standard buffers. Films were reapplied daily following the acid challenge.

### Mineral Preservation

Calcium tablets were treated with simulated saliva for 2 hours, followed by experimental formulas, toothpaste, and distilled water as a negative control. These were then treated with *S. mutans* inoculum in tryptic soy broth for 24 hours, separated from their surrounding solution, and dissolved in acetic acid. Mineral retention was measured using a colorimetric calcium assay (Novus Biologicals Colorimetric Calcium Assay Kit). 50 microliters of sample were treated with the reagent, and absorbance was measured at 570 nm with a spectrophotometer. Raw absorbance values were processed with the linear regression model from concurrently run standards, *y* = 0. 1550*x* + 0. 0045. The limit of detection was established based on the linearity of the 7-point curve (0 to 1.25 mmol/L, *R*^2^ = 0.991). Concentrations were calculated based on the following formula: 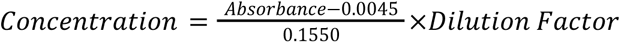

### Gene Expression Analysis (RT-qPCR)

#### Total RNA extraction

Total RNA was extracted from each biofilm sample using the following thermal lysis protocol. Biofilms were physically dislodged from the substrate in cold phosphate-buffered saline (PBS) and harvested by centrifugation at 13,000 x g for 30 seconds. After discarding the supernatant, the bacterial pellet was resuspended in 100 μL of RNase-free water. Thermal lysis was achieved by incubating the suspension in a water bath or dry block heater at 95°C for 10 minutes to disrupt the cell walls and release intracellular nucleic acids. The resulting lysate was immediately cooled on ice for 2 minutes and then centrifuged at 13,000 x g for 5 minutes at 4°C to sediment cellular debris. The RNA-containing supernatant was carefully transferred to a clean, RNase-free microcentrifuge tube. To eliminate residual genomic DNA contamination prior to downstream analysis, the supernatant was treated with DNase I at 37°C for 15 minutes, followed by heat inactivation of the enzyme at 65°C for 10 minutes. It was then run through the Qiagen Rneasy kit following manufacturer instructions. The purified RNA extract was stored at −80°C until RT-qPCR analysis.

While chemical lysis buffers and RNase inhibitors are standard for preserving absolute RNA integrity, this thermal method was selected as a rapid, accessible, and resource-limited extraction approach aligned with the low-cost focus of the global health application, and was supplemented by processing through the Qiagen RNeasy kit. Because the primary objective of this analysis was relative comparative gene expression across treatment groups rather than absolute transcript quantification, any uniform degradation kinetics were normalized across all samples by comparing downstream data to the baseline negative control.

#### cDNA synthesis

Two sets of PCR tubes were prepared for each group. 10 μL of template RNA, 4 μL of 5X Reaction Buffer, 2 μL of dNTP Mix (10 mM), 1 μL of RevertAid Reverse Transcriptase (Thermo Fisher Scientific), and 1 μL of random hexamer primers were added into each tube. Using a 10 μL pipette, each solution was mixed by pipetting. The tubes were centrifuged for 10 sec for the liquid to accumulate in the bottom of the tubes. Then, the tubes were placed in the thermocycler at 25°C for 10 min, 42°C for 60 min, 70°C for 5 min to terminate the reaction and held at 4°C infinitely.

#### Real-time quantitative PCR (RT-qPCR)

To quantify gene expression, 10 μL of SYBR Green Master Mix, 1 μL of forward primer, 1 μL of reverse primer, 7 μL of nuclease-free water, and 1 μL of cDNA were prepared for each reaction. Primers were diluted to a working concentration of 10 pmol/μL. Each reaction mix was vortexed for 5 sec and centrifuged briefly. The RT-qPCR was performed as follows: 1) 95°C for 2 min, 2) 95°C for 15 sec, 3) 60°C for 30 sec, 4) 72°C for 30 sec, and 5) repeat steps 2 to 4 for 40 cycles. For normalization, the 16S rRNA gene was used as an internal housekeeping control. Relative expression levels were determined using the ΔΔCt method. All PCR reagents and primers were purchased from IDT.

**Table 3:**
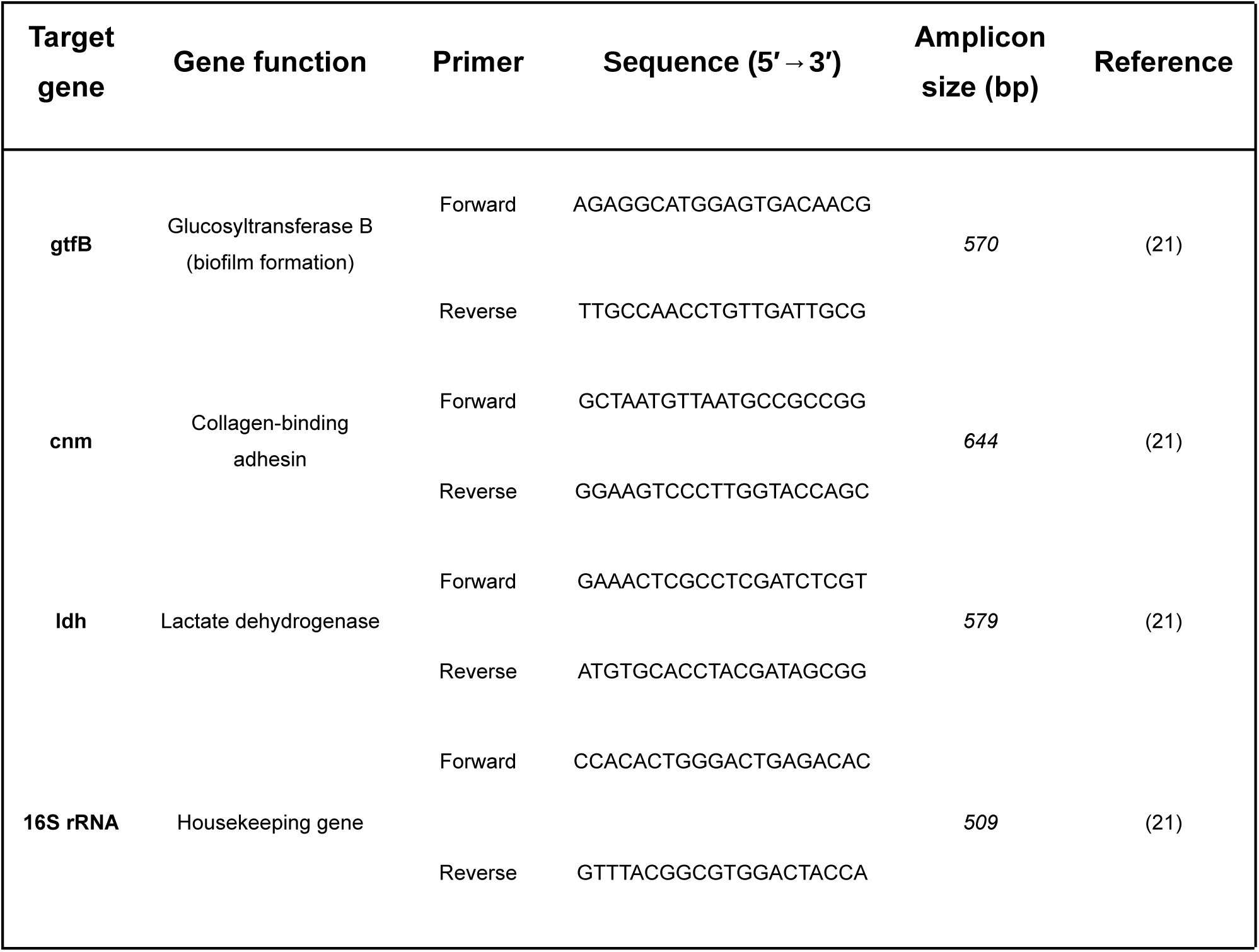
RT-qPCR primer sequences, target genes, and expected amplicon sizes used in this study (21).

### Statistical Analysis

Statistical analyses were performed independently for each experimental phase, with the significance threshold set a priori at alpha = 0.05. Prior to analysis, colony-forming unit (CFU) data were log10-transformed to satisfy the assumption of normality required for parametric testing. For gene expression analysis, raw cycle threshold (Ct) values were normalized to the 16s rRNA housekeeping gene, and relative fold-changes were calculated using the comparative 2^(-ΔΔCt) method. To evaluate differences between the experimental formulations, the toothpaste positive control, and the untreated negative control, a one-way Analysis of Variance (ANOVA) was conducted for each distinct dataset except for pH (log-transformed CFUs, percent cell viability, mean buffered, retained calcium concentration, and relative gene expression). pH was analyzed via a two-way repeated measures ANOVA for the separate values, and mean pH was analyzed via a one-way ANOVA. For all datasets displaying a significant F-statistic, subsequent Tukey’s Honestly Significant Difference (HSD) post hoc tests were applied to isolate specific pairwise differences between groups. All computational analyses and statistical tests were executed using Microsoft Excel.

## RESULTS

The experimental formulations exhibited distinct therapeutic profiles across the microbial, chemical, and genetic evaluation phases. Quantitative analysis revealed significant variations in biofilm burden, cell viability, acid buffering capacity, mineral retention, and virulence gene expression between the targeted treatment groups, the toothpaste positive control, and the untreated negative control.

**Figure 3:**
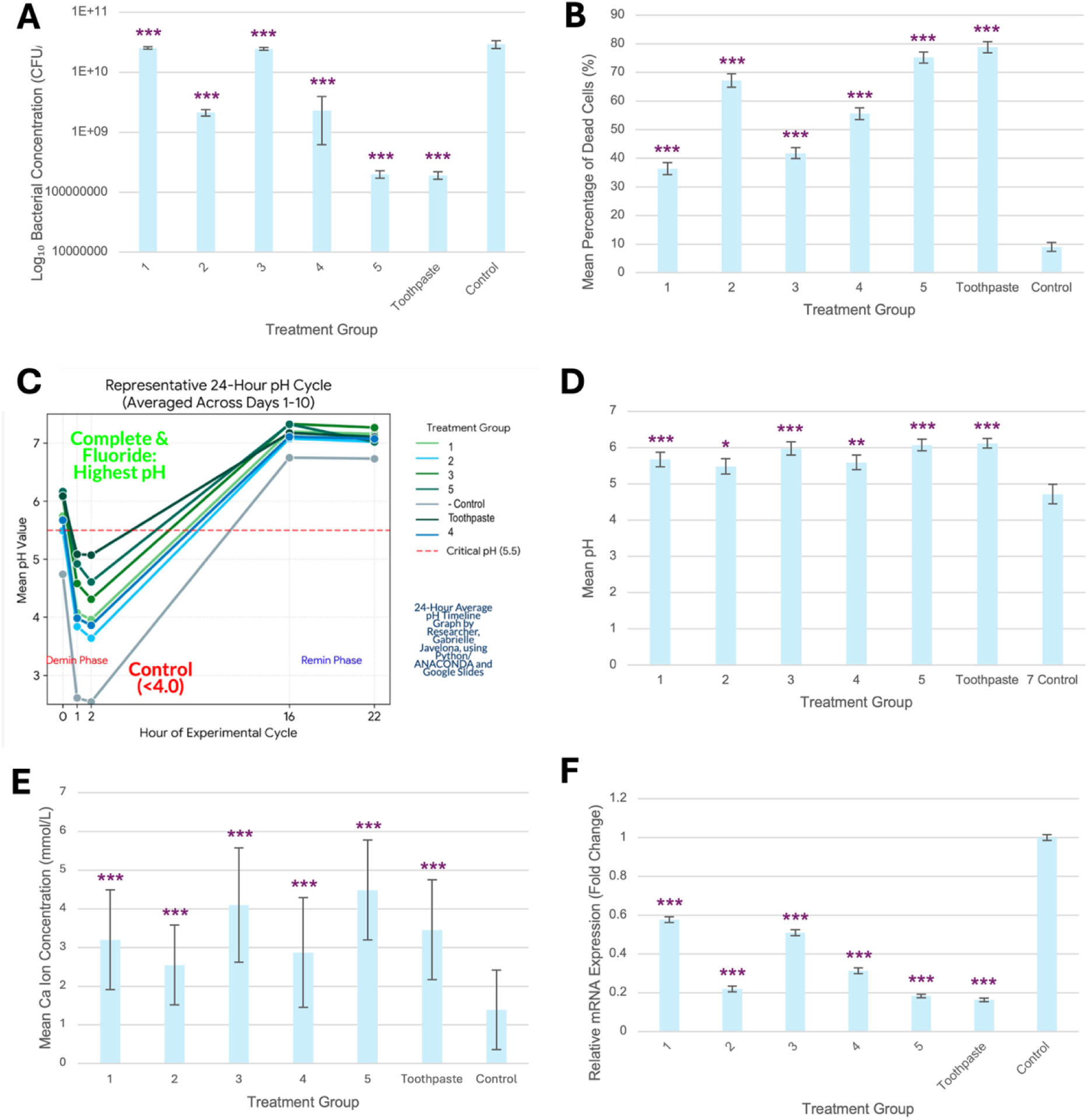
Statistical analysis was performed using one-way ANOVA followed by Tukey’s HSD post-hoc test for panels 3A, 3B, 3D, 3E, and 3F. Panel 3C was analyzed using two-way repeated-measures ANOVA followed by Tukey’s HSD post-hoc test. Data were analyzed using a 95% confidence interval. Statistical significance is indicated as ns (p ≥ 0.05), * (p < 0.05), ** (p < 0.01), *** (p < 0.001), and **** (p < 0.0001). Error bars represent standard deviation. **Figure 3A: Comparison of Log_10_ Bacterial Concentration (CFU/mL) across treatment groups 1-5, Toothpaste, and Control:** Bar graph showing mean ± SD of Log_10_ Bacterial Concentration (n=5). *S. mutans* biofilm was exposed to treatments and controls and cultured on nutrient agar for 24 hours. **Figure 3B: Cell Viability (Amine-Reactive Dye):** Bar graph showing mean ± SD of nonviable cell percentage (%). Cellular viability was quantified using an Invitrogen Amine-Reactive Fixable Dead Cell Stain (n=5). **Figure 3C: 24-Hour pH Cycle:** Line graph showing pH measurements over time: The simulated oral environment was monitored across 10 days using a Beetrode micro-pH electrode (n=5). The red dashed line indicates the critical hydroxyapatite (tooth enamel) dissolving threshold (pH 5.5). **Figure 3D: Mean pH Values:** Cumulative Mean pH Values. Bar graph displaying the cumulative longitudinal mean pH +/- SD for each treatment group, calculated by aggregating the repeated time-course measurements taken across the entire 10-day experimental period (n=5 per group). **Figure 3E: Calcium Ion Concentration:** Bar graph showing mean calcium concentration (mmol/L) ± SD (n=5). The simulated enamel surface Ca^2+^ concentrations were measured using a Calcium Colorimetric Assay Kit. **Figure 3F: Gene Expression (mRNA):** Bar graph showing mean relative mRNA expression ± SD (n=3). RNA was extracted using the Qiagen RNeasy Mini Kit, and cDNA synthesized using the Revertaid cDNA synthesis kit. The target genes (*gtfB*, *cnm*, and *ldh*) were measured with RT-qPCR and normalized to the 16s rRNA housekeeping gene. Relative expression was calculated using the 2^-ΔΔ^Ct method, and the untreated control group served as the calibrator (1.0).

### Biofilm Viability and Disruption (CFU counts)

CFU counts evaluated how well each formula disrupted the formation of *S. mutans* biofilms. After growing *S. mutans* on simulated enamel surfaces (glass beads) for 24 hours, biofilms were dislodged, diluted to 10^-5^, and plated on nutrient agar to quantify surviving colonies.

Significant differences in bacterial concentration were observed between the treatment groups and the negative control (*S. mutans* culture, TSB). The EGCG-DCP-Xyl film (Green Tea, DCP, and Xylitol) showed the greatest biofilm inhibition (1.99 x 10^8^ ± 2.82 x 10^7^ CFU/mL, Table 4). This performance was similar in magnitude to the toothpaste positive control (1.92 x 10^8^ ± 2.83 x 10^7^ CFU/mL; p=0.4238, Table 4), and this demonstrates a significant reduction in biofilm viability compared to the untreated control (2.92 x 10^10^ ± 4.18 x 10^9^ CFU/mL, Table 4; p<0.0001). The Chi-Xyl film performed statistically similarly to the DCP-Xyl film (p=0.1171).

**Table 4:** Mean Colony Forming Units (CFU), mean concentration (CFU/mL), and SD of *S. mutans* per plate when treated with experimental groups and controls. Biofilms were cultured on nutrient agar and quantified via ImageJ.

| Group | Mean CFU | Dilution Multiplier | Mean Concentration (CFU/mL) | SD | Statistical Significance |
| --- | --- | --- | --- | --- | --- |
| 1 | 253.4 | $10^7$ | $2.53 \times 10^{10}$ | $\pm 1.14 \times 10^9$ | <0.001 |
| 2 | 212.8 | $10^6$ | $2.13 \times 10^9$ | $\pm 2.60 \times 10^9$ | <0.001 |
| 3 | 245.2 | $10^7$ | $2.45 \times 10^{10}$ | $\pm 1.34 \times 10^9$ | <0.001 |
| 4 | 228.2 | $10^6$ | $2.28 \times 10^9$ | $\pm 1.66 \times 10^9$ | <0.001 |
| 5 | 198.8 | $10^5$ | $1.99 \times 10^8$ | $\pm 2.82 \times 10^7$ | <0.001 |
| Toothpaste | 192.2 | $10^5$ | $1.92 \times 10^8$ | $\pm 2.83 \times 10^7$ | <0.001 |
| Control | 292.0 | $10^7$ | $2.92 \times 10^{10}$ | $\pm 4.18 \times 10^9$ | 1 |

### Bacterial Killing Efficiency (Live/Dead Fluorescence Staining)

To measure the bactericidal effectiveness of the films, dead fluorescence staining was used. CFU counts measured growth, whereas fluorescence identified the ratio of cells that are actively killed by the treatment. Cells were stained with a fixable red dead cell dye and imaged under a blue laser at 570 nm.

The EGCG-DCP-Xyl film demonstrated the highest mean percentage of non-viable (dead) cells among experimental groups (75% ± 1.92%; p<0.0001, Table 5). This was similar in magnitude to toothpaste (p=0.0999). This was followed by The Chi-Gly film (67% ± 2.39%; p<0.0001, Table 5) and The Gly-Xyl film (56% ± 2.07%; p<0.0001, Table 5). For comparison, the negative control yielded the least *S. mutans* death (9% ± 1.58%, Table 5), while the toothpaste control achieved the most *S. mutans* death. (79% ± 1.92%; p<0.0001, Table 5). These results suggest that the combination of green tea and DCP in the EGCG-DCP-Xyl film is highly effective at inducing cell death in *S. mutans* biofilms.

**Table 5:**
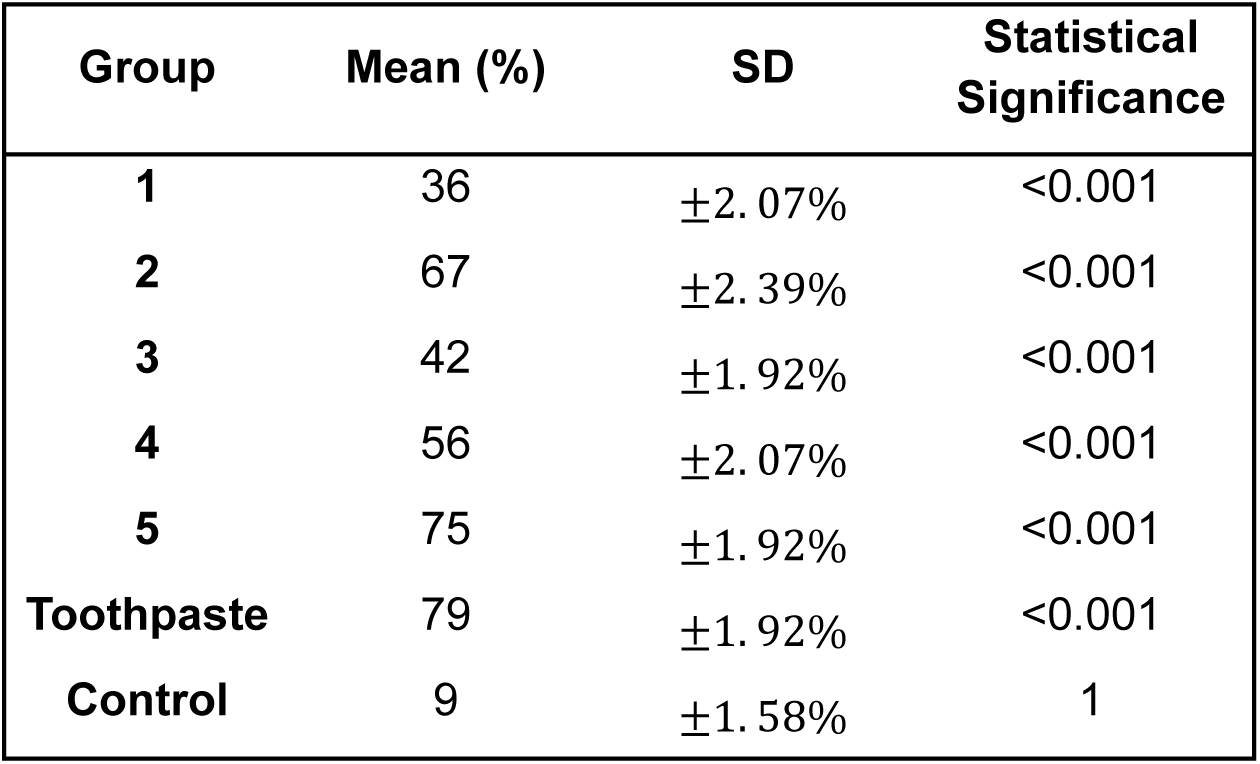
Mean percentages (%) and SD of dead *S. mutans* cells. Percentages were derived from the ratio of dead-cell fluorescence to total cell population.

| Group | Mean (%) | SD | Statistical Significance |
| --- | --- | --- | --- |
| 1 | 36 | $\pm 2.07\%$ | <0.001 |
| 2 | 67 | $\pm 2.39\%$ | <0.001 |
| 3 | 42 | $\pm 1.92\%$ | <0.001 |
| 4 | 56 | $\pm 2.07\%$ | <0.001 |
| 5 | 75 | $\pm 1.92\%$ | <0.001 |
| Toothpaste | 79 | $\pm 1.92\%$ | <0.001 |
| Control | 9 | $\pm 1.58\%$ | 1 |

### pH Buffering and Acid Neutralization (pH Cycling)

The pH cycling experiment simulated the daily acidic and alkaline challenges faced by human enamel in the oral cavity. The primary mechanism of tooth decay is the production of lactic acid by *S. mutans,* which dissolves enamel when the pH drops below 5.5. pH was measured over a 10-day period following acid challenges with Sprite and alkaline challenges with milk.

The untreated control group reached a mean pH well below the critical threshold for enamel demineralization (4.72 ± 2.08, Table 6). All experimental formulas provided a buffering effect that maintained the mean pH above the control levels (p<0.0001). The EGCG-DCP-Xyl film and the DCP-Xyl film provided the strongest buffering (6.07 ± 1.25 and 5.97 ± 1.40, respectively, Table 6) consistently above the critical threshold of 5.5. These values were close to the toothpaste control (6.12 ± 1.03, Table 6), and there was no significant difference in buffering activity between the EGCG-DCP-Xyl film and toothpaste (p=0.60) suggesting that the film ingredient formulas can successfully neutralize the lactic acid produced by *S. mutans*.

**Table 6:** Mean pH levels across 10 days and SD for each experimental group. Acid and alkaline challenges were administered, and pH monitored with a probe at set time intervals.

| Group | Mean pH | SD | Statistical Significance |
| --- | --- | --- | --- |
| 1 | 5.67 | 1.57 | 0.0054 |
| 2 | 5.48 | 1.64 | 0.034 |
| 3 | 5.97 | 1.40 | 0.0002 |
| 4 | 5.59 | 1.56 | 0.0109 |
| 5 | 6.07 | 1.25 | <0.0001 |
| Toothpaste | 6.12 | 1.03 | <0.0001 |
| Control | 4.72 | 2.08 | 1 |

### Mineral Retention and Preservation (Calcium Assay)

After being treated with the different film formulas, the calcium tablets simulating tooth enamel were separated from their surrounding fluid and dissolved in acetic acid. Following this, a calcium colorimetric assay measured mineral retention, the ability of the film ingredient combinations to prevent the leaching of calcium from the simulated enamel surface during acid challenges.

Higher absorbance at 570 nm indicated a greater concentration of retained calcium ions. The EGCG-DCP-Xyl film exhibited the highest concentration among the experimental groups (4.487 mmol/L ± 1.290; p<0.0001, Table 7) outperforming The Chi-Xyl film (3.197 mmol/L ± 1.290; p<0.0001, Table 7) and the DCP-Xyl film (4.100 mmol/L ± 1.484; p=0.0141, Table 7). This data suggested that the inclusion of dicalcium phosphate (DCP) and green tea in the EGCG-DCP-Xyl film significantly enhances the mineral stability of the simulated enamel surface compared to the other film variations. There was no significant difference in mineral retention between Formulas 2 and 4 (p=0.066), and between The Chi-Xyl film and toothpaste (p=0.0877)

**Table 7:** Absorbance and concentration of calcium in samples at a wavelength of 570 nm. The absorbance of calcium ions released quantified from the in vitro simulated enamel surface via a spectrometer.

| Group | Mean | Mean Concentration | SD | Statistical Significance |
| --- | --- | --- | --- | --- |
| 1 | 0.50 | 3.197 | 1.290 | <0.001 |
| 2 | 0.40 | 2.552 | 1.032 | <0.001 |
| 3 | 0.64 | 4.100 | 1.484 | <0.001 |
| 4 | 0.45 | 2.874 | 1.419 | <0.001 |
| 5 | 0.70 | 4.487 | 1.290 | <0.001 |
| Toothpaste | 0.54 | 3.455 | 1.290 | <0.001 |
| Control | 0.22 | 1.390 | 1.032 | 1 |

The high calcium concentrations in combination with the pH stability maintained by the EGCG-DCP-Xyl film suggest that a direct mineral scaffold with nHA and DCP might exceed fluoride’s mineral preservation potential, although more robust testing (Vickers microhardness, SEM imaging) is required.

### Gene Expression of Virulence Factors (RT-qPCR)

To understand the molecular mechanism behind these observations, RT-qPCR was used to analyze the expression of the *gtfB* (biofilm formation), *cnm* (virulence), and *ldh* (lactate dehydrogenase) genes. Ct values were normalized to the 16srRNA housekeeping gene. In The EGCG-DCP-Xyl film, there was a noticeable downregulation in the expression of *gtfB*, *cnm*, and *ldh* (0.183 ± 0.015, 0.220 ± 0.020, and 0.230 ± 0.020 respectively, Table 8) compared to the control group (p<0.0001). In *gtfB* expression, there was no significant difference between the EGCG-DCP-Xyl film and toothpaste (p=0.9337), and between Formulas 2 and 5 (p=0.5001). In *cnm* expression, there was no significant difference between the EGCG-DCP-Xyl film and toothpaste (p=0.9889), and between Formulas 1 and 3 (p=0.466, Table 8). In *ldh* expression, there was no significant difference between Formulas 2 and 4 (p=0.2903), and between the EGCG-DCP-Xyl film and toothpaste (p=0.8937). This reduction in gene expression correlates with the reduced CFU counts and higher pH levels observed in previous phases, indicating that the ingredients in the film interfere with the bacteria’s ability to produce sticky biofilms and acidic waste.

**Table 8:** Relative fold expression of *gtfB*, *cnm*, and *ldh* genes in *S. mutans*: Mean expression values represent the fold-change relative to the control group (normalized to 1.000) using the 2^-ΔΔ^Ct method.

| Group | *gtfB* Mean Expression | *cnm* Mean Expression | *ldh* Mean Expression | *gtfB* SD | *cnm* SD | *ldh* SD | Statistical Significance |
| --- | --- | --- | --- | --- | --- | --- | --- |
| 1 | 0.577 | **0.607** | 0.580 | 0.025 | **0.025** | 0.030 | <0.001 |
| 2 | 0.220 | **0.317** | 0.350 | 0.026 | **0.025** | 0.020 | <0.001 |
| 3 | 0.510 | **0.570** | 0.540 | 0.026 | **0.020** | 0.026 | <0.001 |
| 4 | 0.313 | **0.390** | 0.397 | 0.025 | **0.020** | 0.025 | <0.001 |
| 5 | 0.183 | **0.220** | 0.230 | 0.015 | **0.020** | 0.020 | <0.001 |
| Toothpaste | 0.163 | **0.207** | 0.207 | 0.015 | **0.015** | 0.015 | <0.001 |
| Control | 1.000 | **1.000** | 1.000 | 0.026 | **0.030** | 0.030 | 1 |

The suppression observed in the *gtfB* and *cnm* genes with the EGCG-DCP-Xyl film aligns with the claim that these pathways are critical for the structural integrity of oral microbial communities (8). Therefore, all the ingredient combinations, but particularly the EGCG-DCP-Xyl film, might prevent the virulence of *S. mutans*, the main bacteria behind dental decay.

## DISCUSSION

This study investigated the effects of different natural ingredient formulas in a film delivery system on the markers of tooth decay, for daily treatment in water-scarce areas. It was hypothesized that an experimental formulation incorporating nano-hydroxyapatite (nHA), xylitol, dicalcium phosphate (DCP), and epigallocatechin gallate (EGCG) would successfully reduce Streptococcus mutans abundance and optimize calcium preservation within a simulated enamel environment, performing comparably to a standard toothpaste positive control. To evaluate this, *S. mutans* biofilm accumulation was quantified via colony-forming unit (CFU) reduction, while cellular viability and metabolic lactic acid suppression were monitored through fluorescence staining and targeted gene-expression analysis by RT-qPCR. Mineral retention of the different formulas was measured through a calcium assay.

Future experimental approaches could utilize Fourier-transform infrared spectroscopy (FTIR) to characterize chemical bonding shifts within the film matrix, alongside molecular docking simulations to map the precise binding affinities between the active compounds and targeted bacterial surface proteins. (9, 10, 11).

In this study, the EGCG-DCP-Xyl film appeared to significantly impact both the metabolic activity of *S. mutans* and the rate of mineral dissolution. EGCG was associated with reduced expression of gtfB, while xylitol was associated with reduced expression of ldh and improved pH stability. Simultaneously, nano-hydroxyapatite and DCP provided calcium and phosphate ions to promote remineralization. By targeting bacterial adhesion, acidogenesis, and mineral loss simultaneously, the combined formulation may have produced complementary effects across these pathways.

This could be explored in the future using Scanning Electron Microscopy (SEM) to visualize surface morphology changes, or Energy-Dispersive X-ray Spectroscopy (EDS) to map the exact calcium-to-phosphate ratio (Ca/P) following film application.

In *S. mutans* biofilm models with CFU counts, bacterial concentration was lowest in the EGCG-DCP-Xyl film group (1.99 x 10^8^ ± 2.82 x 10^7^ CFU/mL; p<0.0001, Table 4), similar in magnitude to commercial toothpaste (12). Since the expression of the *gtfB* gene was significantly reduced (0.183 ± 0.015; p<0.0001, Table 8), the reduction in CFU counts likely stems from a disruption in the production of glucosyltransferases (Figure 1, Source 13). Lower gtfB expression limits extracellular glucan synthesis required for mature biofilm formation, whereas decreased ldh expression reduces lactic acid production responsible for enamel demineralization.

This also suggests that the EGCG-DCP-Xyl film does not only prevent attachment but may compromise the structural integrity of the bacterial cell wall, potentially through the polyphenolic action of EGCG (Figure 1). This may induce oxidative stress and membrane damage in Gram-positive cocci like *S. mutans*, as described by Reygaert et al. (14).

The 10-day pH cycling assay also supports the film’s effects on pH stability, where the EGCG-DCP-Xyl film maintained a mean pH consistently above the critical demineralization threshold of 5.5 (6.07 ± 1.25, Table 6). This correlates with the reduction in expression of the *ldh* gene (0.230 ± 0.020, Table 8), suggesting that the film suppresses the conversion of carbohydrates into lactic acid via xylitol, as displayed in Arunakul et al. (Figure 1, Source 9).

Although this research was designed to be as comprehensive as possible, there were some limitations to this project. First, the pH cycling model was conducted in a simplified in vitro environment using a static broth medium. This does not account for the constant flushing action of saliva, which can mechanically rinse bacterial acids away. In the future, a constant-depth film fermentor (CDFF) could simulate the continuous flow of saliva and forces of chewing, providing more realistic conditions to test how well the film adheres (5).

Secondly, qPCR for gene expression yielded a general idea of the metabolic state of *S. mutans*, but it does not account for the translational level of protein synthesis. Although a reduction of expression was recorded in all three of the relevant virulence genes, it was impossible to determine if the actual production of each associated protein was reduced in the same magnitude. Flow cytometry or a Western blot assay could address this in the future. Finally, the CFU counts and Live/Dead Fluorescence on biofilms were grown on uniform surfaces, which may not replicate the more complex architecture (pits, cavities) of human teeth. Each formula’s performance could be assessed against different bacteria like Fusobacterium nucleatum and Candida albicans to simulate the complex oral microbiome in the future, as well.

Future experimentation could also determine the exact pathways by which these formulas affect the oral microbiome. The film formulas’ modification of the oral biofilm genetics could also be measured over extended periods to prevent resurgence of acidogenic bacteria like *S. mutans* after application. Flow cytometry could confirm the protein expression of *S. mutans* after treatment, and advanced microscopy could be used to confirm the exact mineral density of the enamel surface after treatment. Specifically, the in vitro simulated model using glass beads was an isolated environment, so analysis of human enamel using Vickers microhardness or Scanning Electron Microscopy (SEM) could confirm surface morphology repairs. Additionally, goats or other animal models could assess the long-term biocompatibility of the film with gingival (gum) tissue before human clinical trials.

Long term trials modeling chronic oral conditions, such as early-stage “white spot” lesions, could also be conducted in the future. A clinical analysis of patients using these film delivery systems could confirm a correlation between formula application and a decrease in cavity progression, as well as a relative dentin abrasivity (RDA) of 250 or less.

To prevent *S. mutans* from developing resistance to the formula ingredients, the most optimal concentrations of ingredients could be optimized further, or the different formula alternatives (DCP-Xyl film for mineral loss prevention or Chi-Gly film for antimicrobial activity) could be used instead.

In summary, the rampant issue of dental decay affects 27.5% of the global population to this day (1). A large percentage of these people lack access to safe water, and the global water bankruptcy declared by UNICEF makes water-free, affordable daily hygiene methods for teeth like this edible film infinitely more important.

## ACKNOWLEDGMENTS

The authors would like to express their sincere gratitude to the Silver River Museum and Environmental Education Center (a program of Marion County Public Schools) for providing $1210.66 in funding and crucial access to Biosafety Level 2 (BSL-2) laboratory facilities, without which this research would not have been possible. Special thanks are extended to Mrs. Christina Davis for her exceptional guidance, technical mentorship, and unwavering support throughout the experimentation process. Additionally, the authors would like to thank Mrs. Erin Benavides for her invaluable coordination, assistance, and dedication to facilitating this science research initiative.

## AUTHOR CONTRIBUTIONS

Gabrielle Javelona: Conceptualization, Investigation, Data Curation, Formal Analysis, Visualization, Writing – Original Draft.

Jacob Merle: Supervision, Project Administration, Writing – Review & Editing.

